# Intermittent Nicotine Access is as Effective as Continuous Access in Promoting Nicotine Seeking and Taking in Rats

**DOI:** 10.1101/2023.10.26.563876

**Authors:** Hajer E Algallal, Vincent Jacquemet, Anne-Noël Samaha

**Affiliations:** Biomedical Sciences, Faculty of Medicine, Université de Montréal; Institute of Biomedical Engineering, Faculty of Medicine, Université de Montréal; Department of Pharmacology and Physiology, Faculty of Medicine, Université de Montréal; Neural Signaling and Circuitry Research Group (SNC), Faculty of Medicine, Université de Montréal; Center for Interdisciplinary Research on the Brain and Learning (CIRCA), Université de Montréal

**Keywords:** Nicotine self-administration, Long Access, Intermittent Access, Progressive Ratio Schedule, Conditioned stimulus, Reinstatement

## Abstract

Nicotine is a principal psychoactive agent in tobacco, contributing to tobacco’s addictive potential. Preclinical studies on the effects of voluntary nicotine intake typically use self-administration procedures that provide continuous nicotine access during each self-administration session. However, many smokers consume cigarettes intermittently rather than continuously throughout each day. For drugs including cocaine and opioids, research in laboratory rats shows that intermittent intake can be more effective than continuous intake in producing patterns of drug use relevant to addiction. We asked, therefore, how intermittent versus continuous nicotine self-administration influences nicotine seeking and taking behaviours. Female and male rats had continuous (i.e., Long Access; LgA, 6 h/day) or intermittent (IntA; 12 min ON, 60 min OFF, for 6 h/day) access to intravenous nicotine (15 µg/kg/infusion), for 12 daily sessions. We then assessed intake, responding for nicotine under a progressive ratio schedule of drug reinforcement and cue- and nicotine-induced reinstatement of drug seeking. We also estimated nicotine pharmacokinetic parameters during LgA and IntA self-administration. Overall, LgA rats took twice more nicotine than did IntA rats, yielding more sustained increases in estimated brain concentrations of the drug. However, the two groups showed similar motivation to seek and take nicotine, as measured using reinstatement and progressive ratio procedures, respectively. Thus, intermittent nicotine use is just as effective as continuous use in producing addiction-relevant behaviours, despite significantly less nicotine exposure. This has implications for modeling nicotine self-administration patterns in human smokers and resulting effects on brain and behaviour.

## Introduction

Tobacco consumption is a global health problem. All forms of tobacco are harmful, and cigarette smoking is the most common form of tobacco use worldwide (World Health Organisation, 2020). In Western Europe, Canada and the United States, tobacco consumption accounts for one in five deaths (Danaei et al., 2009; Hughes, 2016; Lim et al., 2012) making cigarette smoking a leading and essentially preventable cause of death. Effective pharmacological and behavioural interventions exist but no treatment works for all smokers (Livingstone-Banks et al., 2022). Even with evidence-based treatments, only 10-30% of smokers achieve sustained abstinence (Clinical Practice Guideline Treating Tobacco Use and Dependence 2008 Update Panel, Liaisons, and Staff, 2008). Six months to a year after a given attempt to quit, only 3-5% of smokers will still be abstinent (Hughes et al., 2004) and quitting smoking successfully can require as many as 30 or more quit attempts (Chaiton et al., 2016).

Cigarette smoke contains several pharmacologically active compounds, but nicotine is the main compound thought to underlie the reinforcing and addictive properties of cigarette smoke. The advent of new anti-smoking treatment options rests on a better understanding of the mechanisms underlying nicotine’s reinforcing and potentially addictive effects. Animal models are useful to achieve this goal. Studies using voluntary nicotine self-administration procedures show nicotine seeking and taking behaviour across species, including dogs (Risner & Goldberg, 1983), rodents (Corrigall & Coen, 1989), and non-human primates (Goldberg et al., 1981; Sannerud et al., 1994). To date, nicotine self-administration procedures have typically provided near-continuous nicotine access during each session lasting 6 h (Paterson & Markou, 2004), 12 h (Kenny & Markou, 2006) or even 23 h (LeSage et al., 2002) per day. Such procedures arguably model chain-smoking, where cigarettes are consumed continuously for many hours each day. Yet, in many smokers, cigarette and nicotine intake is intermittent, both within and between days. Daily smokers can have peaks and troughs in the number of cigarettes they smoke over a day, as well as repeated cigarette smoking in short time windows (Zhai et al., 2022). Non-daily smokers also show variation in smoking rates and in the number of smoking bouts on the days when they do smoke (Braznell et al., 2022; Li et al., 2022; Shiffman et al., 2014). Food, caffeine and alcohol consumption, work, social setting, cigarette availability and mood all contribute to the intermittency of cigarette smoking (Shiffman et al., 2014; Stennett et al., 2018).

Many studies using cocaine have compared the effects of continuous (long access; LgA) versus intermittent (IntA) drug access and found that IntA is uniquely effective in promoting behavioural changes relevant to addiction (Allain et al., 2015; Kawa et al., 2019; Samaha et al., 2021). LgA provides drug continuously, typically during 4-6-h self-administration sessions (Ahmed & Koob, 1999), whereas IntA involves signalled periods of drug availability and unavailability during each session (Zimmer et al., 2011, 2012). Despite resulting in less cocaine intake compared to LgA, IntA promotes long-lasting psychomotor sensitization (Algallal et al., 2020; Allain et al., 2017; Carr et al., 2020), persistent increases in the motivation to take drug (Algallal et al., 2020; Allain & Samaha, 2019; Calipari et al., 2014; James et al., 2019), and robust cue-induced reinstatement of drug seeking (Gueye et al., 2019; James et al., 2019; Kawa et al., 2016, 2019; Singer et al., 2018). Other preclinical studies have also shown that intermittency is important for the effects of other drugs, including heroin (O’Neal et al., 2020) and fentanyl (Fragale et al., 2019). Thus, beyond how much drug is taken, the temporal pattern of drug use can be decisive in predicting outcome.

How intermittent versus continuous access may influence nicotine self-administration behaviour is largely unknown. Tapia et al (2022) gave rats intermittent nicotine access and found that they reliably self-administered the drug and showed more cue-induced nicotine seeking at 7 vs. 1 day of abstinence. These rats also showed alterations in the dorsal striatum proteome and evidence of increased binding and function of nicotinic acetylcholine receptors (Tapia et al., 2022). Thus, IntA procedures support nicotine self-administration behaviour and produce neurobiological changes. Our objective here was to directly compare the effects of IntA vs. LgA nicotine on several drug-seeking and drug-taking behaviours relevant to addiction. To this end, we compared drug consumption patterns, estimated nicotine pharmacokinetic profiles, responding for nicotine under a progressive ratio schedule of reinforcement and extinction conditions, and cue- and nicotine-induced reinstatement of nicotine-seeking behaviour.

## Materials and Methods

See the Supplement for information on subjects, apparatus, surgeries, drugs, self-administration training and data analysis. The animal care committee of the Université de Montréal approved all experimental procedures, and these complied with the guidelines of the Canadian Council on Animal Care.

### IntA and LgA sessions

Figure 1 shows the sequence of experimental events. After intravenous catheter implantation and self-administration training, female (150-175 g) and male (225-250 g) Wistar rats (Charles River Laboratories, Saint Constant, QC, Canada) were divided into two groups. One group received 6-h IntA sessions (n = 10), and the other group received 6-h LgA sessions (n = 10). The rats were assigned to each group such that groups were matched on the mean number of self-administered nicotine injections and lever presses on the last two training sessions, and the mean number of days to meet acquisition criteria. All rats then self-administered nicotine infusions (0.015 mg/kg/infusion, delivered over 5 s; MP Biomedicals, Solon, Ohio, US) paired with a tone-light conditioned stimulus (CS). The rats received 1 session/day, every other day for a total of 12 sessions. Each IntA session consisted of five 12-min nicotine ON periods, where nicotine was available under FR3 without a timeout period (save for each 5-s injection). ON periods, which were signalled by active and inactive lever insertion, were intercalated with 60-min, nicotine OFF periods where levers were retracted and nicotine was unavailable. Nicotine t_1/2_ in rat plasma and brain is ∼60 min (Hwa Jung et al., 2001; Kyerematen et al., 1988; Kyerematen & Vesell, 1991; Schechter & Jellinek, 1975). As such, nicotine concentrations would be expected to decrease during each 60-min OFF period and to peak again during the next ON period. This would produce the spiking pattern of drug concentrations in the brain characteristic of IntA (Algallal et al. 2020; Allain et al. 2017; Allain et al. 2018; Tapia et al. 2022; Zimmer et al. 2012). During LgA sessions, nicotine was available continuously under FR3, except for a 20-s timeout period following each self-administered injection. Central nicotine concentrations were estimated using self-administration time series data from the 12th LgA and IntA sessions (see Supplement). Following the 12th IntA or LgA session, rats were kept in their home cages for 4 days before progressive ratio testing.

**Figure 1.**
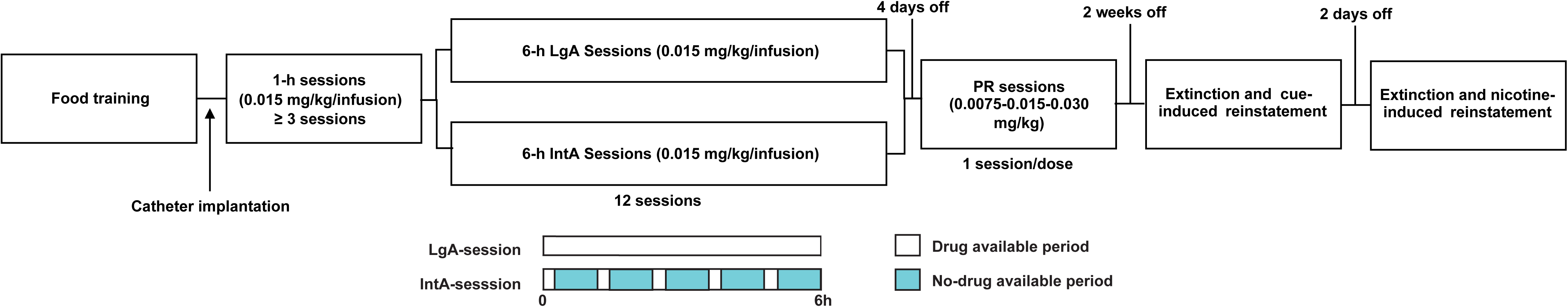
The sequence of experimental events. Female and male rats were trained to press a lever for food pellets. Subsequently, catheters were implanted into the jugular vein. After recovery from surgery, rats were allowed to self-administer nicotine during 1-h sessions, and then assigned to self-administer nicotine during 6-h Long Access sessions (LgA) or 6-h Intermittent Access sessions (IntA), for 12 sessions (1/day). Four days following the last session, we measured responding under a progressive ratio schedule of nicotine reinforcement (PR). Then, three weeks following the last LgA or IntA session, we assessed cue and nicotine-induced reinstatement of nicotine-seeking behavior.

### Progressive ratio (PR)

Four days following the last IntA or LgA session, we assessed incentive motivation for nicotine by determining responding for nicotine (0.0075, 0.015 and 0.030 mg/kg/injection, in counterbalanced order, 1 session/dose) under a PR schedule (Richardson & Roberts, 1996).

### Abstinence, extinction and reinstatement tests

Three weeks after the last LgA/IntA session, all rats received two 6-h extinction sessions (1/day on consecutive days). During each session, both levers were present throughout, and pressing the levers produced neither nicotine nor the CS. Immediately after the second extinction session, a 1-h CS-induced reinstatement test was initiated. The session started with a response-noncontingent presentation of the nicotine CS. Thereafter, active lever presses produced the CS, under FR3. No nicotine was delivered during this test. Two days later, each rat received two more 6-h extinction sessions (1/day). Immediately after each extinction session, rats received a s.c. injection of saline or nicotine [0.15 mg/kg in a volume of 1 ml/kg (Le et al. 2006; Shaham et al. 1997)], in a within-subjects design and in counterbalanced order. A 1-h nicotine (or saline)-induced reinstatement test was then initiated. During reinstatement tests, pressing on either lever had no programmed consequences.

## Results

### Modeled plasma nicotine concentrations under IntA vs. LgA

We estimated plasma nicotine pharmacokinetics on the 12th self-administration session (Fig. 2). Fig. 2A shows average estimated nicotine concentrations over a 10-h period in each group, encompassing the 6-h self-administration session and up to 4 h after. IntA produced a spiking pattern of nicotine concentrations, with peaks and troughs (Figure 2A), whereas LgA produced more sustained increases in nicotine concentrations [Figure 2A; see also (Tapia et al., 2022)]. IntA and LgA rats reached similar nicotine Cmax values, both as estimated across the 6-h self-administration session (respectively, 21.81 ng/ml, and 27.37 ng/ml, Fig. 2B; *p* > 0.05) and within the first 12-min of the session (respectively, 9.56 ng/ml, and 8.72 ng/ml, Fig. 2C; *p* > 0.05). Within the first 12-min of the session (corresponding to the first nicotine ON period in the IntA rats), IntA and LgA rats also showed similar estimated nicotine Tmax values (respectively, 9.66 min, and 10.17 min, Fig. 2D; *p >* 0.05). However, across all timepoints over a 24-h period, IntA rats had significantly lower estimated area under the curve (AUC) values for plasma nicotine compared to LgA rats (main effect of Access; F_1,11_ = 9.60, *p* = 0.01; main effect of Time; F_3,33_ = 61.26, *p* < 0.0001; Time X Access interaction effect; F_3,33_ = 5.04, *p* = 0.005; Fig. 2D). Thus, compared to LgA rats, IntA rats had repeated spikes in nicotine concentrations as well as significantly lower overall nicotine exposure.

**Figure 2.**
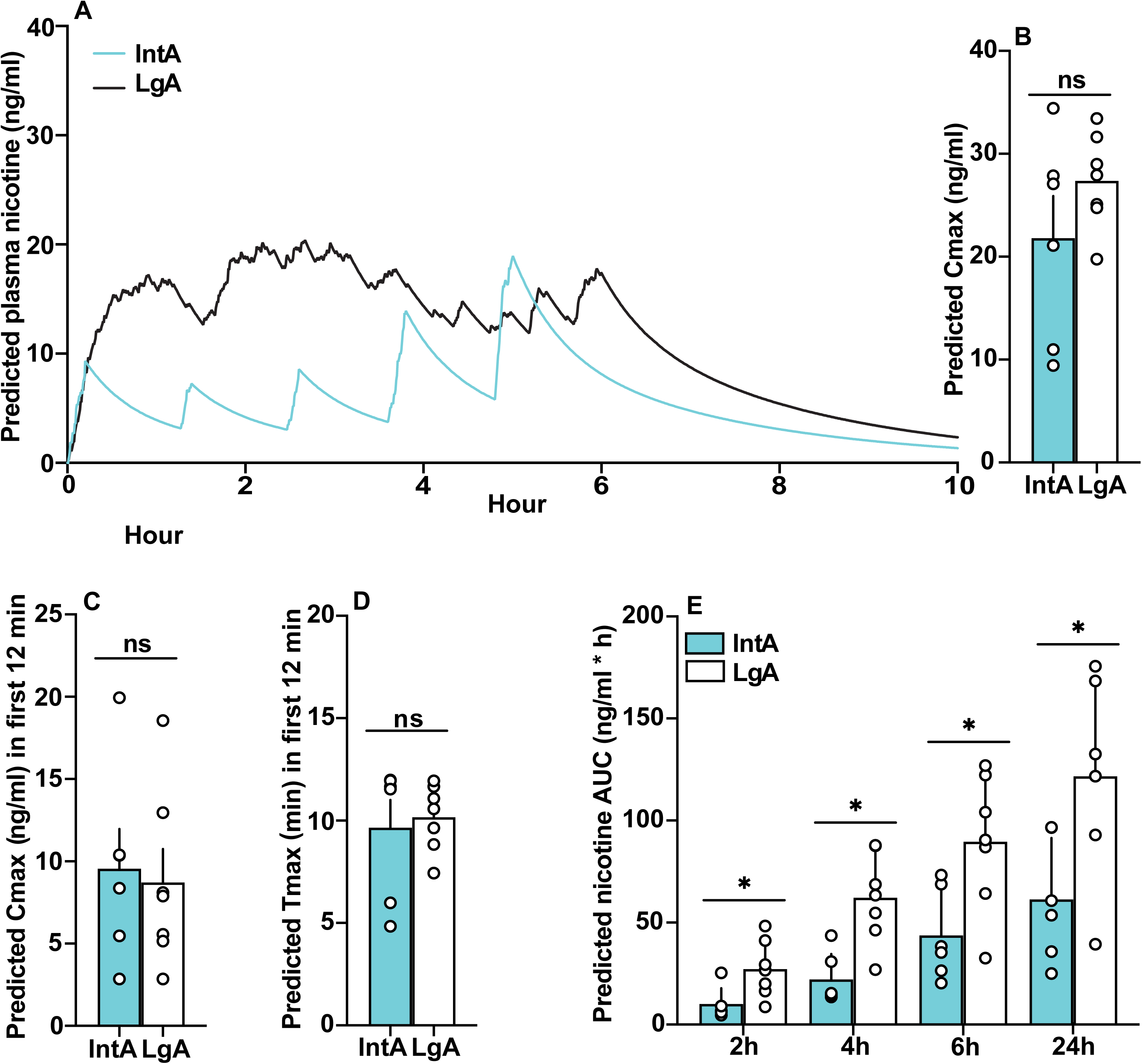
Intermittent (IntA) and Long access (LgA) nicotine self-administration produce different estimated pharmacokinetic profiles. Plasma nicotine concentrations were modeled on the 12^th^ day of IntA or LgA nicotine self-administration in male rats (see text). (**A)** Predicted time course of changes in plasma nicotine concentrations with IntA versus LgA self-administration. LgA produced sustained elevations in estimated nicotine concentrations. Under IntA, nicotine concentrations would instead follow a spiking pattern. (**B**) Predicted peak plasma nicotine concentrations (Cmax) across the 6-h self-administration session. (**C**) Predicted peak plasma nicotine concentrations in the first 12 min of the session, corresponding to the first nicotine ON period for IntA rats. (**D**) Predicted time to reach peak plasma nicotine concentrations (Tmax) in the first 12 min of the session. (**E**) Predicted area under the curve (AUC) for plasma nicotine over time. * *p* < 0.05. n = 10 males.

### Nicotine self-administration behaviour during IntA vs. LgA sessions

Lever-pressing behaviour during nicotine self-administration training, IntA/LgA, progressive ratio and reinstatement sessions was similar across the sexes and they were pooled for analysis. Figure 3 shows nicotine self-administration behaviour in IntA and LgA rats. Figs. 3A-B show representative patterns of nicotine intake in each group. LgA rats typically self-administered nicotine in more sustained fashion across each 6-h session. In contrast, IntA rats necessarily took nicotine in intermittent bouts. Across groups, rats significantly discriminated between the two levers, pressing more on the active vs. inactive lever (Fig. 3C; Lever; F_1,18_ = 85.75, *p* < 0.001). Across groups, lever pressing behaviour also varied as function of Session (F_11,198_ = 3.54, *p* < 0.001; no other comparisons were significant). LgA rats pressed more on both levers as compared to IntA rats (Fig. 3C; Access; F_1,18_ = 12.29, *p* = 0.003; Lever type X Access; F_1,18_ = 9.37, *p* < 0.01; Active lever: Access; F_1,18_ = 13.02, *p* = 0.002; Session; F_11,189_ = 1.93, *p* = 0.03; Inactive lever; main effect of Access; F_11,18_ = 10.17, *p* = 0.005; Session; F_11,189_ = 3.46, *p* = 0.0002). Accordingly, LgA rats also took more nicotine infusions than IntA rats did (Fig. 3D; Access; F_1,18_ = 13.54, *p* = 0.001. No other comparisons were statistically significant). There were group differences in cumulative nicotine intake over the 12 self-administration sessions (Fig. 3E; *p* = 0.004), such that LgA rats took on average 2.6-fold more nicotine than IntA rats did.

**Figure 3.**
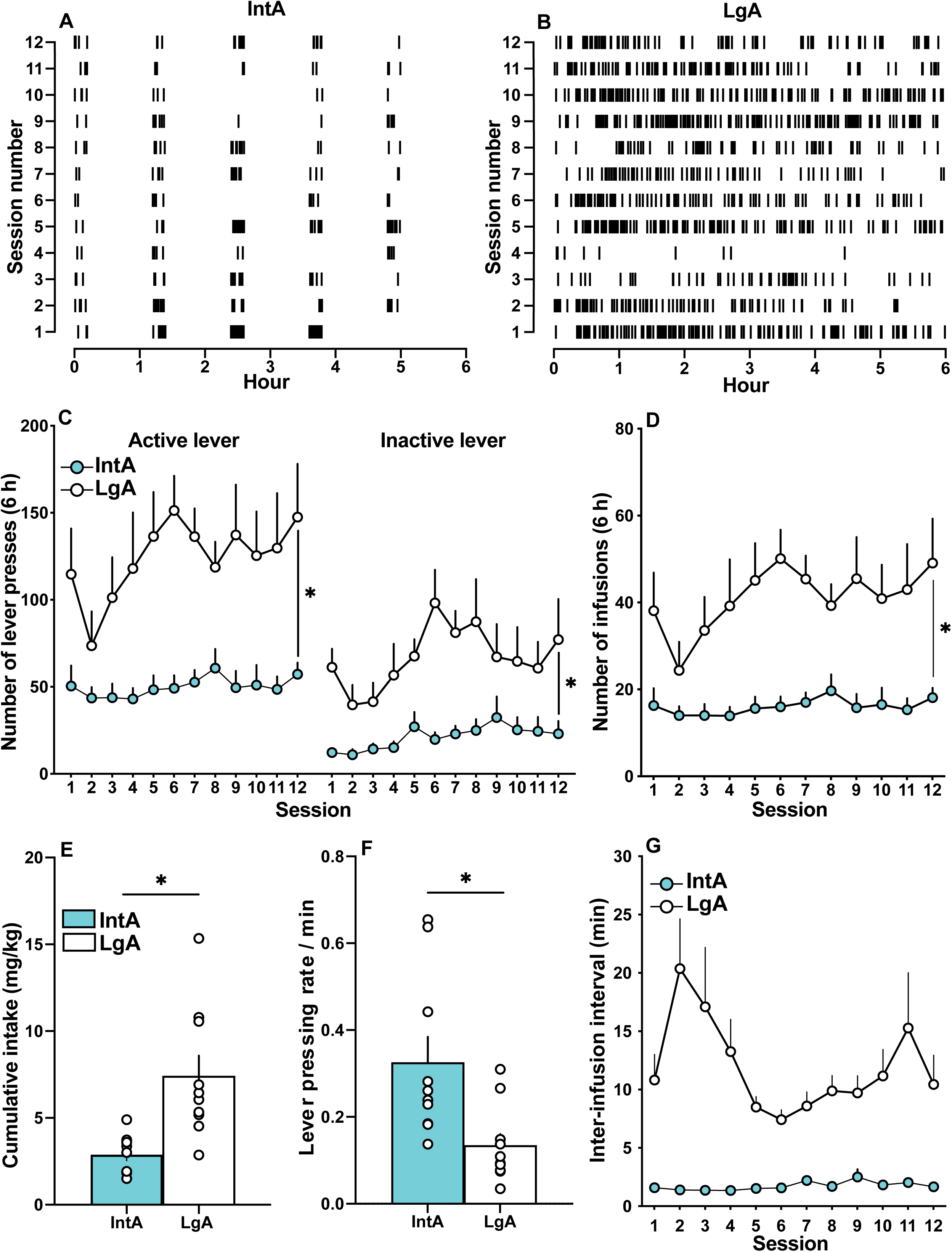
Intermittent (IntA) and Long access (LgA) produce distinct patterns of nicotine self-administration behaviour. Patterns of nicotine intake in (**A**) a representative IntA rat and (**B**) a representative LgA rat, across the 6 h of each of the 12 self-administration sessions. Each tick mark represents one self-administered infusion. (**C**) Both IntA and LgA groups pressed significantly more on the active versus inactive lever, and lever pressing was highest in the LgA rats. (**D**) Across sessions, LgA rats took significantly more nicotine infusions than did IntA rats. (**E**) Cumulative nicotine intake was greatest in LgA rats. (**F**) IntA rats responded for nicotine at higher rates than did LgA rats (calculated as the ratio of total number of infusions obtained/session over the total number of active lever presses/session, averaged over the last 4 self-administration sessions). (**G**) Compared to LgA rats, IntA rats self-administered nicotine infusions at significantly shorter latencies*. *p* < 0.05, Data are mean ± SEM. n = 7 females, 13 males.

We analysed group differences in the rate of responding for nicotine, over the last 4 self-administration sessions (Fig. 3F). IntA rats responded at higher rates for nicotine compared to LgA rats (Fig. 3F; *p* = 0.009). Accordingly, IntA rats also self-administered nicotine at shorter inter-infusion intervals (Fig. 3G; Access; F_1,18_ = 27.45, *p* < 0.0001). In summary, LgA rats took more nicotine than IntA rats did, but IntA rats responded at higher rates for the drug.

With cocaine, IntA promotes increased drug intake at the beginning of each drug-available period of the session, and this loading effect can sensitise over sessions (Algallal et al., 2020; Allain et al., 2018; Kawa & Robinson, 2019). We examined this with nicotine here. Supplementary Figure 1 shows average number of nicotine infusions taken during each 3-min bin of the 12-min nicotine ON periods, over the 1^st^, 6^th^ and 12^th^ IntA sessions. In contrast to findings with cocaine (Algallal et al., 2020; Allain et al., 2018; Kawa & Robinson, 2019), IntA rats did not load up on nicotine at the beginning of each ON period. Instead, they took nicotine at relatively constant rates across each ON period (Supplementary Figure 1; Time (min); F_3,32_ = 0.38, *p >* 0.05), and this pattern of intake remained unchanged across the 12 IntA sessions (Supplementary Figure 1; effect of Session; F_1.7,57.4_ = 0.56, *p >* 0.05). We also analyzed the latency to self-administer a nicotine infusion after the start of each ON period for IntA rats (Supplementary Figure 1B) and after the start of each session for LgA rats (Supplementary Figure 1C). IntA rats showed a range of latencies, and the most common latencies were between 20 and 60 s (Supplementary Figure 1B). In LgA rats, the most common latency before initiating self-administration was between 120 and 140 s (Supplementary Figure 1C). In summary, during periods of signaled nicotine availability, IntA rats rapidly initiate self-administration and take nicotine at a steady rate, both within and across self-administration sessions. LgA rats initiate self-administration after longer latencies, and they then also take nicotine at steady rates.

### Progressive Ratio Responding

Four days after the last IntA or LgA session, rats received PR tests (0.0075-0.030 mg/kg/infusion nicotine; Fig. 4). Across groups, rats pressed more on the active vs. inactive lever (main effect of Lever F_1,18_ = 46.89, *p* < 0.001; Fig. 4A), and they responded more for higher nicotine doses, particularly on the active lever (Dose; F_2,36_ = 9.02, *p* = 0.001; Lever type X Dose; F_2,36_ = 5.18, *p* = ; Fig. 4A; no other comparisons were significant). Across groups, the number of earned infusions was dose-dependent (main effect of Dose; F_2,36_ = 4.99, *p* = 0.01; Fig. 4B). Across nicotine doses, IntA and LgA rats earned a similar number of nicotine infusions (Dose x Access and Access, all *p* values > 0.05; Fig. 4B). Sessions lasted longer with increasing nicotine dose (Dose; F_2,36_ = 5.35, *p* = 0.01; Fig. 4C), and there were no group differences in this effect (All *p* values > 0.05). Thus, rats with a history of intermittent versus continuous nicotine self-administration responded similarly for nicotine under PR.

**Figure 4.**
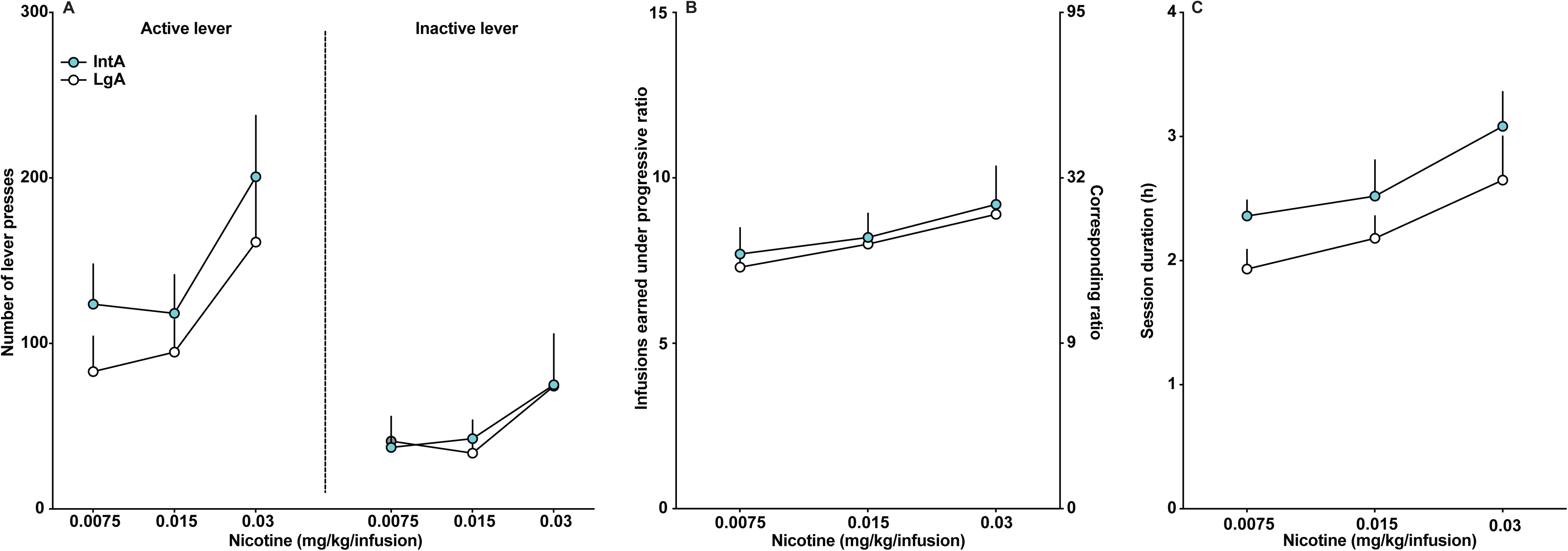
Intermittent access (IntA) and Long access (LgA) rats show similar responding for nicotine during progressive ratio tests. (**A**) Both IntA and LgA groups pressed significantly more on the active versus inactive lever, and lever pressing was similar in the two groups. (**B**) The two groups earned a similar number of nicotine infusions, across a range of nicotine doses. (**C**) Session duration was similar between IntA and LgA rats. Data are mean ± SEM. n = 7 females, 13 males.

### Cue- and nicotine-induced reinstatement

Rats decreased responding over the 6-h extinction session and there were no group differences in this effect (Fig. 5A; Time (h); F_5,90_ = 17.53, *p* < 0.001; no other comparisons were significant). Across groups, rats pressed more on the previously nicotine-associated (active) lever during the CS-induced reinstatement test than during the last hour of extinction (Fig. 5B; Session type; F_1,18_ = 37.40, *p* < 0.0001; Figs. 5B; Lever type; F_1,18_ = 30.13, *p* < 0.0001; Lever type X Session type; F_1,18_ = 35.99, *p* < 0.0001). There were no significant group differences in responding (Figs. 5B; Access; *p* > 0.05). Thus, IntA and LgA rats showed similar levels of CS-induced reinstatement.

**Figure 5.**
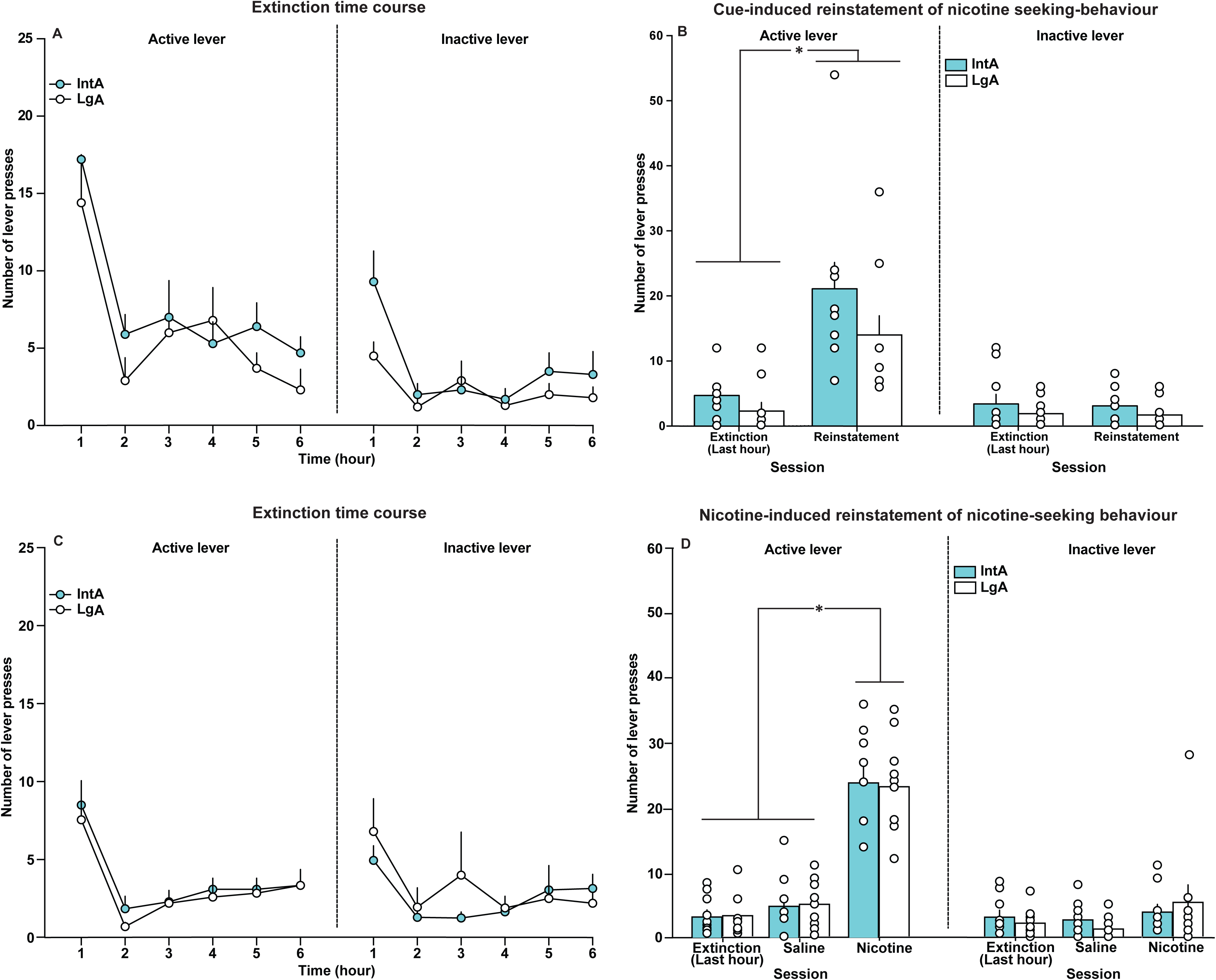
Rats with a history of self-administering nicotine under Intermittent (IntA) or Long (LgA) access conditions show similar responding for nicotine under extinction sessions and similar levels of cue- and nicotine-induced reinstatement of drug-seeking behaviour. (**A**) Active and inactive lever pressing behaviour during extinction was similar in IntA and LgA rats. (**B**) IntA and LgA rats showed significant cue-induced reinstatement of extinguished nicotine-seeking behaviour, with no group differences. (**C**) Active and inactive lever pressing behaviour during extinction was similar in IntA and LgA rats. (**D**) A priming injection of nicotine (0.15 mg/kg, s.c.) triggered similar levels of reinstatement of nicotine-seeking behaviour in the two groups. *\*p* < 0.05, Main effect of session type. Data are mean ± SEM. n = 7 females, 13 males.

Two days after the CS-induced reinstatement test, rats received two more extinction/reinstatement sessions, where each 6-h extinction session was immediately followed by a 1-h saline- or nicotine-induced reinstatement test. Responding on the two extinction sessions was similar and Fig. 5C shows lever-pressing behaviour averaged across the two sessions. Across groups, rats decreased responding during extinction training (Fig. 5C; Time (h); F_5,90_ = 15.49, *p* < 0.001; no other comparisons were significant). Fig. 5D shows responding during reinstatement tests vs. extinction. Across groups, a priming injection of nicotine increased responding on the active lever relative to both saline and extinction conditions (Fig. 5D; Session type; F_2,36_ = 67.04, *p* < 0.001; Lever type; F_1,18_ = 72.87, *p* < 0.001; Lever type x Session type; F_2,36_ = 71.87, *p* < 0.001). There were no group differences in responding (Fig. 5D; Access; *p* > 0.05). In summary, IntA rats and LgA rats showed similar extinction responding, and cue- and nicotine-induced reinstatement of drug-seeking behaviour.

### Correlations between IntA/LgA self-administration behaviour and later responding during PR or reinstatement tests

Cumulative nicotine intake, inter-infusion interval or lever pressing rate during prior IntA/LgA sessions generally did not significantly predict responding during PR tests [all *p* values > .003; Supplementary table 1; see also (Algallal et al., 2020; Allain et al., 2017, 2021)] or active lever pressing during cue- or nicotine-induced reinstatement tests (all *p* values > .003; Supplementary table 2). These observations extend prior work conducted with cocaine (Allain et al., 2018, 2021; Kawa & Robinson, 2019; Oleson & Roberts, 2009; Zimmer et al., 2012) and show that measures of nicotine consumption can be dissociable from later appetitive responding for the drug.

## Discussion

We compared the effects of intermittent (IntA) versus continuous (LgA) nicotine self-administration on nicotine-seeking and -taking behaviours in rats. We report two main findings. First, IntA and LgA produced different patterns of nicotine self-administration behaviour and predicted pharmacokinetic profiles. LgA rats took more nicotine infusions across sessions as compared to IntA rats. However, during periods where nicotine was available, IntA rats initiated self-administration faster and also responded at higher rates for nicotine compared to LgA rats. LgA self-administration also produced more sustained elevations in estimated plasma nicotine concentrations, whereas nicotine concentrations followed a spiking pattern with IntA [see also (Tapia et al., 2022)]. Second, though IntA rats had significantly less nicotine exposure than did LgA rats, the two groups showed similar responding for the drug under progressive ratio and extinction conditions and similar cue- and nicotine-induced reinstatement of drug seeking. Thus, intermittent nicotine use is as effective as continuous use in producing addiction-relevant behaviours, but with markedly less total nicotine exposure.

### IntA and LgA produce similar behavioural outcomes despite differences in total nicotine exposure

Compared to LgA rats, IntA rats took two-fold less nicotine overall, yet the two groups later showed similar motivation to seek and take the drug. Thus, less frequent nicotine use and reduced levels of cumulative drug intake (as modeled by IntA) can be as effective as more continuous and heavier intake (as modeled by LgA) in producing addiction-relevant behaviours. To the extent that our findings can be generalized to humans, an intermittent pattern of nicotine/cigarette use may be sufficient to promote tobacco use disorder. Extending recent work (Tapia et al., 2022), our IntA procedure can be used to identify the behavioural and neurobiological mechanisms that sustain intermittent nicotine intake. This is especially important as some treatments that can be effective in heavier/daily smokers can be ineffective in intermittent smokers (Shiffman et al., 2020).

### How IntA to nicotine may increase motivation for nicotine despite reduced drug exposure

Intermittent peaks in brain nicotine concentrations evoked by IntA may promote sensitization-related neuroadaptations within dopamine systems, thus increasing incentive motivation for the drug, as suggested by studies using IntA cocaine self-administration (Allain et al., 2021; Calipari et al., 2013; Calipari & Jones, 2014; Carr et al., 2020; Kawa et al., 2019). Sensitization-related plasticity has yet to be examined with IntA nicotine self-administration. However, intermittent, experimenter-delivered i.v. infusions of nicotine promote such plasticity, albeit with doses higher than used here (Lenoir et al., 2013; Samaha et al., 2005).

Learning processes may also contribute. With IntA, the lever is a discriminative cue associated with a high rate of reinforcement. This would result in high levels of Pavlovian excitation being conditioned to the lever after IntA versus LgA training (Beasley et al., 2022), thus producing increased reward-seeking and reward-taking behaviours despite less exposure to drug reward with IntA. Our IntA rats indeed responded for nicotine at higher rates during self-administration sessions and also took nicotine at shorter intervals compared to LgA rats (Fig. 3F-G). However, response rate did not significantly predict responding for nicotine during progressive ratio or reinstatement tests. Thus, while IntA rats showed increased response rates, this may not have contributed to subsequent nicotine-seeking and -taking behaviours.

IntA may also promote the ability of nicotine to increase the incentive motivational effects of nicotine-associated cues, thereby enhancing drug pursuit when such cues are present (Caggiula et al., 2009; Donny et al., 2003; Goldberg et al., 1981). However, a nicotine conditioned stimulus produced similar reinstatement behaviour in our IntA and LgA rats. Nonetheless, we did not characterize the many psychological and behavioural effects through which drug-associated cues contribute to drug use. For instance, cue-induced cocaine relapse is greater following IntA vs. LgA experience (Nicolas et al., 2019), suggesting that IntA can promote the incentive motivational properties of drug-associated cues.

Whatever the underlying mechanisms, our findings may also suggest that the nicotine threshold for addiction is lower than previously thought, such that despite reduced total nicotine exposure, lighter/intermittent nicotine consumption can still produce the forms of plasticity that promote further nicotine use. Thus, beyond the amount of nicotine taken, the intermittency of use may contribute to sustaining nicotine self-administration.

### IntA and smoking profiles in humans

Our IntA procedure may be useful to model intermittent nicotine exposure relevant to humans. Studies of smoking patterns using ecological momentary assessment and related methods suggest that some smokers have peaks and troughs in the number of cigarettes smoked across the day, as well as variation in the number of daily smoking bouts (Braznell et al., 2022; Li et al., 2022; Shiffman et al., 2014). In addition, 20 to 40% of current smokers are estimated to be light and intermittent smokers [i.e., smoking < 10 cigarettes/day and not smoking every day (Reyes-Guzman et al., 2017)]. Light/intermittent smoking is not simply a transition stage, as it does not necessarily lead to heavier smoking (Caldeira et al., 2012; Levy et al., 2009). This highlights the importance of developing animal models of light/intermittent nicotine intake.

### Methodological Considerations

We could not consider sex as a biological variable in data analysis because we lost many female rats to catheter failure. At least some effects of nicotine/tobacco smoking can be different in women versus men (Jarvis et al., 2013). Some preclinical studies in rats report sex differences in nicotine self-administration behaviour (Chaudhri et al., 2005; Donny et al., 2000; Grebenstein et al., 2013; Pogun et al., 2017; Sanchez et al., 2014) while others report no or only minor sex differences (Feltenstein et al., 2012; Swalve et al., 2016). Future work can extend the present findings across the sexes. We used 15 µg/kg/infusion, because it supports high rates of self-administration behaviour under fixed ratio schedules of reinforcement (Corrigall & Coen, 1989; Matta et al., 2007). In addition, our predicted peak nicotine concentrations (Fig. 2B) are within the smoking-relevant range of around 20 ng/ml (Lawson et al., 1998). However, 15 µg/kg/infusion is higher than that extracted by humans with each puff on a cigarette [1 to 3 µg/kg nicotine/puff, and 10 to 30 µg/kg/cigarette (Matta et al., 2007)]. Nicotine pharmacokinetics differ between rats and humans making it difficult to translate doses across species. However, rats will self-administer lower, ‘puff-sized’ nicotine doses i.v (Donny et al., 1995; Sorge & Clarke, 2009; Watkins et al., 1999). Thus, it will be worthwhile to extend the present findings to additional nicotine doses.

## Conclusions

We compared nicotine self-administration behaviour in rats given IntA vs. LgA to the drug. LgA rats took twice more nicotine than did IntA rats and the two groups also showed different temporal patterns of nicotine self-administration and pharmacokinetic profiles. However, the two groups later showed similar responding for nicotine during progressive ratio, extinction and reinstatement tests. Thus, lighter/intermittent nicotine use, which is relevant to a significant proportion of cigarette smokers, can be modelled effectively in laboratory rats [see also (Tapia et al., 2022)]. Moreover, despite producing less nicotine exposure overall, intermittent nicotine self-administration can be just as effective as more continuous self-administration in promoting features relevant to tobacco use disorder.

## Acknowledgements

We thank Dr. Paul Clarke for reading and commenting an earlier version of this manuscript.

## Author Contributions

HEA and ANS designed the research. HEA performed all experiments and analyzed all data with guidance from ANS. VJ modelled nicotine concentrations. HEA and ANS wrote the article with revisions from VJ.

## References

Ahmed, S. H., & Koob, G. F. (1999). Long-lasting increase in the set point for cocaine self-administration after escalation in rats. Psychopharmacology, 146(3), Article 3. 10.1007/s002130051121

Algallal, H., Allain, F., Ndiaye, N. A., & Samaha, A.-N. (2020). Sex differences in cocaine self-administration behaviour under long access versus intermittent access conditions. Addiction Biology, 25(5), Article 5. 10.1111/adb.12809

Allain, F., Bouayad-Gervais, K., & Samaha, A.-N. (2018). High and escalating levels of cocaine intake are dissociable from subsequent incentive motivation for the drug in rats. Psychopharmacology, 235(1), Article 1. 10.1007/s00213-017-4773-8

Allain, F., Delignat-Lavaud, B., Beaudoin, M.-P., Jacquemet, V., Robinson, T. E., Trudeau, L.-E., & Samaha, A.-N. (2021). Amphetamine maintenance therapy during intermittent cocaine self-administration in rats attenuates psychomotor and dopamine sensitization and reduces addiction-like behavior. Neuropsychopharmacology: Official Publication of the American College of Neuropsychopharmacology, 46(2), Article 2. 10.1038/s41386-020-0773-1

Allain, F., Minogianis, E.-A., Roberts, D. C. S., & Samaha, A.-N. (2015). How fast and how often: The pharmacokinetics of drug use are decisive in addiction. Neuroscience & Biobehavioral Reviews, 56, 166–179. 10.1016/j.neubiorev.2015.06.012

Allain, F., Roberts, D. C. S., Lévesque, D., & Samaha, A.-N. (2017). Intermittent intake of rapid cocaine injections promotes robust psychomotor sensitization, increased incentive motivation for the drug and mGlu2/3 receptor dysregulation. Neuropharmacology, 117, 227–237. 10.1016/j.neuropharm.2017.01.026

Allain, F., & Samaha, A.-N. (2019). Revisiting long-access versus short-access cocaine self-administration in rats: Intermittent intake promotes addiction symptoms independent of session length. Addiction Biology, 24(4), Article 4. 10.1111/adb.12629

Beasley, M. M., Gunawan, T., Tunstall, B. J., & Kearns, D. N. (2022). Intermittent access training produces greater motivation for a non-drug reinforcer than long access training. Learning & Behavior, 50(4), Article 4. 10.3758/s13420-022-00512-w

Braznell, S., Van Den Akker, A., Metcalfe, C., Taylor, G. M. J., & Hartmann-Boyce, J. (2022). Critical appraisal of interventional clinical trials assessing heated tobacco products: A systematic review. Tobacco Control, tobaccocontrol-2022–057522. 10.1136/tc-2022-057522

Caggiula, A. R., Donny, E. C., Palmatier, M. I., Liu, X., Chaudhri, N., & Sved, A. F. (2009). CHAPTER 6: THE ROLE OF NICOTINE IN SMOKING: A DUAL-REINFORCEMENT MODEL. Nebraska Symposium on Motivation. Nebraska Symposium on Motivation, 55, 91–109.

Caldeira, K. M., O’Grady, K. E., Garnier-Dykstra, L. M., Vincent, K. B., Pickworth, W. B., & Arria, A. M. (2012). Cigarette Smoking Among College Students: Longitudinal Trajectories and Health Outcomes. Nicotine & Tobacco Research, 14(7), Article 7. 10.1093/ntr/nts131

Calipari, E. S., Ferris, M. J., Siciliano, C. A., Zimmer, B. A., & Jones, S. R. (2014). Intermittent Cocaine Self-Administration Produces Sensitization of Stimulant Effects at the Dopamine Transporter. The Journal of Pharmacology and Experimental Therapeutics, 349(2), Article 2. 10.1124/jpet.114.212993

Calipari, E. S., Ferris, M. J., Zimmer, B. A., Roberts, D. C., & Jones, S. R. (2013). Temporal Pattern of Cocaine Intake Determines Tolerance vs Sensitization of Cocaine Effects at the Dopamine Transporter. Neuropsychopharmacology, 38(12), Article 12. 10.1038/npp.2013.136

Calipari, E. S., & Jones, S. R. (2014). Sensitized nucleus accumbens dopamine terminal responses to methylphenidate and dopamine transporter releasers after intermittent-access self-administration. Neuropharmacology, 82, 1–10. 10.1016/j.neuropharm.2014.02.021

Carr, C. C., Ferrario, C. R., & Robinson, T. E. (2020). Intermittent access cocaine self-administration produces psychomotor sensitization: Effects of withdrawal, sex and cross-sensitization. Psychopharmacology, 237(6), Article 6. 10.1007/s00213-020-05500-4

Chaiton, M., Diemert, L., Cohen, J. E., Bondy, S. J., Selby, P., Philipneri, A., & Schwartz, R. (2016). Estimating the number of quit attempts it takes to quit smoking successfully in a longitudinal cohort of smokers. BMJ Open, 6(6), Article 6. 10.1136/bmjopen-2016-011045

Chaudhri, N., Caggiula, A. R., Donny, E. C., Booth, S., Gharib, M. A., Craven, L. A., Allen, S. S., Sved, A. F., & Perkins, K. A. (2005). Sex differences in the contribution of nicotine and nonpharmacological stimuli to nicotine self-administration in rats. Psychopharmacology, 180(2), Article 2. 10.1007/s00213-005-2152-3

Clinical Practice Guideline Treating Tobacco Use and Dependence 2008 Update Panel, Liaisons, and Staff. (2008). A clinical practice guideline for treating tobacco use and dependence: 2008 update. A U.S. Public Health Service report. American Journal of Preventive Medicine, 35(2), Article 2. 10.1016/j.amepre.2008.04.009

Corrigall, W. A., & Coen, K. M. (1989). Nicotine maintains robust self-administration in rats on a limited-access schedule. Psychopharmacology, 99(4), Article 4. 10.1007/BF00589894

Danaei, G., Ding, E. L., Mozaffarian, D., Taylor, B., Rehm, J., Murray, C. J. L., & Ezzati, M. (2009). The preventable causes of death in the United States: Comparative risk assessment of dietary, lifestyle, and metabolic risk factors. PLoS Medicine, 6(4), Article 4. 10.1371/journal.pmed.1000058

Donny, E. C., Caggiula, A. R., Knopf, S., & Brown, C. (1995). Nicotine self-administration in rats. Psychopharmacology, 122(4), Article 4. 10.1007/BF02246272

Donny, E. C., Caggiula, A. R., Rowell, P. P., Gharib, M. A., Maldovan, V., Booth, S., Mielke, M. M., Hoffman, A., & McCallum, S. (2000). Nicotine self-administration in rats: Estrous cycle effects, sex differences and nicotinic receptor binding. Psychopharmacology, 151(4), Article 4. 10.1007/s002130000497

Donny, E. C., Chaudhri, N., Caggiula, A. R., Evans-Martin, F. F., Booth, S., Gharib, M. A., Clements, L. A., & Sved, A. F. (2003). Operant responding for a visual reinforcer in rats is enhanced by noncontingent nicotine: Implications for nicotine self-administration and reinforcement. Psychopharmacology, 169(1), Article 1. 10.1007/s00213-003-1473-3

Feltenstein, M. W., Ghee, S. M., & See, R. E. (2012). Nicotine self-administration and reinstatement of nicotine-seeking in male and female rats. Drug and Alcohol Dependence, 121(3), Article 3. 10.1016/j.drugalcdep.2011.09.001

Fragale, J. E., Pantazis, C. B., James, M. H., & Aston-Jones, G. (2019). The role of orexin-1 receptor signaling in demand for the opioid fentanyl. Neuropsychopharmacology, 44(10), Article 10. 10.1038/s41386-019-0420-x

Goldberg, S. R., Spealman, R. D., & Goldberg, D. M. (1981). Persistent behavior at high rates maintained by intravenous self-administration of nicotine. Science (New York, N.Y.), 214(4520), Article 4520. 10.1126/science.7291998

Grebenstein, P., Burroughs, D., Zhang, Y., & LeSage, M. G. (2013). Sex differences in nicotine self-administration in rats during progressive unit dose reduction: Implications for nicotine regulation policy. Pharmacology, Biochemistry, and Behavior, 114–115, 70–81. 10.1016/j.pbb.2013.10.020

Gueye, A. B., Allain, F., & Samaha, A.-N. (2019). Intermittent intake of rapid cocaine injections promotes the risk of relapse and increases mesocorticolimbic BDNF levels during abstinence. Neuropsychopharmacology, 44(6), Article 6. 10.1038/s41386-018-0249-8

Hughes, J. R. (2016). National Institutes of Health Funding for Tobacco Versus Harm From Tobacco. Nicotine & Tobacco Research, 18(5), Article 5. 10.1093/ntr/ntv137

Hughes, J. R., Keely, J., & Naud, S. (2004). Shape of the relapse curve and long-term abstinence among untreated smokers. Addiction (Abingdon, England), 99(1), Article 1. 10.1111/j.1360-0443.2004.00540.x

Hwa Jung, B., Chul Chung, B., Chung, S. J., & Shim, C. K. (2001). Different pharmacokinetics of nicotine following intravenous administration of nicotine base and nicotine hydrogen tartarate in rats. Journal of Controlled Release: Official Journal of the Controlled Release Society, 77(3), Article 3. 10.1016/s0168-3659(01)00452-7

James, M. H., Stopper, C. M., Zimmer, B. A., Koll, N. E., Bowrey, H. E., & Aston-Jones, G. (2019). Increased Number and Activity of a Lateral Subpopulation of Hypothalamic Orexin/Hypocretin Neurons Underlies the Expression of an Addicted State in Rats. Biological Psychiatry, 85(11), Article 11. 10.1016/j.biopsych.2018.07.022

Jarvis, M. J., Cohen, J. E., Delnevo, C. D., & Giovino, G. A. (2013). Dispelling myths about gender differences in smoking cessation: Population data from the USA, Canada and Britain. Tobacco Control, 22(5), Article 5. 10.1136/tobaccocontrol-2011-050279

Kawa, A. B., Bentzley, B. S., & Robinson, T. E. (2016). Less is more: Prolonged intermittent access cocaine self-administration produces incentive-sensitization and addiction-like behavior. Psychopharmacology, 233(19–20), Article 19–20. 10.1007/s00213-016-4393-8

Kawa, A. B., & Robinson, T. E. (2019). Sex differences in incentive-sensitization produced by intermittent access cocaine self-administration. Psychopharmacology, 236(2), Article 2. 10.1007/s00213-018-5091-5

Kawa, A. B., Valenta, A. C., Kennedy, R. T., & Robinson, T. E. (2019). Incentive and dopamine sensitization produced by intermittent but not long access cocaine self-administration. European Journal of Neuroscience, 50(4), Article 4. 10.1111/ejn.14418

Kenny, P. J., & Markou, A. (2006). Nicotine self-administration acutely activates brain reward systems and induces a long-lasting increase in reward sensitivity. Neuropsychopharmacology: Official Publication of the American College of Neuropsychopharmacology, 31(6), Article 6. 10.1038/sj.npp.1300905

Kyerematen, G. A., Taylor, L. H., deBethizy, J. D., & Vesell, E. S. (1988). Pharmacokinetics of nicotine and 12 metabolites in the rat. Application of a new radiometric high performance liquid chromatography assay. Drug Metabolism and Disposition: The Biological Fate of Chemicals, 16(1), Article 1.

Kyerematen, G. A., & Vesell, E. S. (1991). Metabolism of nicotine. Drug Metabolism Reviews, 23(1–2), Article 1–2. 10.3109/03602539109029754

Lawson, G. M., Hurt, R. D., Dale, L. C., Offord, K. P., Croghan, I. T., Schroeder, D. R., & Jiang, N. S. (1998). Application of serum nicotine and plasma cotinine concentrations to assessment of nicotine replacement in light, moderate, and heavy smokers undergoing transdermal therapy. Journal of Clinical Pharmacology, 38(6), Article 6. 10.1002/j.1552-4604.1998.tb05787.x

Lenoir, M., Tang, J. S., Woods, A. S., & Kiyatkin, E. A. (2013). Rapid Sensitization of Physiological, Neuronal, and Locomotor Effects of Nicotine: Critical Role of Peripheral Drug Actions. The Journal of Neuroscience, 33(24), Article 24. 10.1523/JNEUROSCI.4940-12.2013

LeSage, M. G., Keyler, D. E., Shoeman, D., Raphael, D., Collins, G., & Pentel, P. R. (2002). Continuous nicotine infusion reduces nicotine self-administration in rats with 23-h/day access to nicotine. Pharmacology, Biochemistry, and Behavior, 72(1–2), Article 1–2. 10.1016/s0091-3057(01)00775-4

Levy, D. E., Biener, L., & Rigotti, N. A. (2009). The natural history of light smokers: A population-based cohort study. Nicotine & Tobacco Research: Official Journal of the Society for Research on Nicotine and Tobacco, 11(2), Article 2. 10.1093/ntr/ntp011

Li, X., Anderson, S. J., Shiffman, S., & Zhang, B. (2022). Time-varying coefficient cumulative gap time models for intensive longitudinal ecological momentary assessment data with missingness. Journal of Applied Statistics, 49(2), Article 2. 10.1080/02664763.2020.1815676

Lim, S. S., Vos, T., Flaxman, A. D., Danaei, G., Shibuya, K., Adair-Rohani, H., Amann, M., Anderson, H. R., Andrews, K. G., Aryee, M., Atkinson, C., Bacchus, L. J., Bahalim, A. N., Balakrishnan, K., Balmes, J., Barker-Collo, S., Baxter, A., Bell, M. L., Blore, J. D.,… Memish, Z. A. (2012). A comparative risk assessment of burden of disease and injury attributable to 67 risk factors and risk factor clusters in 21 regions, 1990-2010: A systematic analysis for the Global Burden of Disease Study 2010. Lancet (London, England), 380(9859), Article 9859. 10.1016/S0140-6736(12)61766-8

Livingstone-Banks, J., Lindson, N., Hartmann-Boyce, J., & Aveyard, P. (2022). Effects of interventions to combat tobacco addiction: Cochrane update of 2019 and 2020 reviews. Addiction (Abingdon, England), 117(6), Article 6. 10.1111/add.15769

Matta, S. G., Balfour, D. J., Benowitz, N. L., Boyd, R. T., Buccafusco, J. J., Caggiula, A. R., Craig, C. R., Collins, A. C., Damaj, M. I., Donny, E. C., Gardiner, P. S., Grady, S. R., Heberlein, U., Leonard, S. S., Levin, E. D., Lukas, R. J., Markou, A., Marks, M. J., McCallum, S. E.,… Zirger, J. M. (2007). Guidelines on nicotine dose selection for in vivo research. Psychopharmacology, 190(3), Article 3. 10.1007/s00213-006-0441-0

Nicolas, C., Russell, T. I., Pierce, A. F., Maldera, S., Holley, A., You, Z.-B., McCarthy, M. M., Shaham, Y., & Ikemoto, S. (2019). Incubation of Cocaine Craving After Intermittent-Access Self-administration: Sex Differences and Estrous Cycle. Biological Psychiatry, 85(11), Article 11. 10.1016/j.biopsych.2019.01.015

Oleson, E. B., & Roberts, D. C. S. (2009). Behavioral economic assessment of price and cocaine consumption following self-administration histories that produce escalation of either final ratios or intake. Neuropsychopharmacology: Official Publication of the American College of Neuropsychopharmacology, 34(3), Article 3. 10.1038/npp.2008.195

O’Neal, T. J., Nooney, M. N., Thien, K., & Ferguson, S. M. (2020). Chemogenetic modulation of accumbens direct or indirect pathways bidirectionally alters reinstatement of heroin-seeking in high-but not low-risk rats. Neuropsychopharmacology: Official Publication of the American College of Neuropsychopharmacology, 45(8), Article 8. 10.1038/s41386-019-0571-9

Paterson, N. E., & Markou, A. (2004). Prolonged nicotine dependence associated with extended access to nicotine self-administration in rats. Psychopharmacology, 173(1–2), Article 1–2. 10.1007/s00213-003-1692-7

Pogun, S., Yararbas, G., Nesil, T., & Kanit, L. (2017). Sex differences in nicotine preference. Journal of Neuroscience Research, 95(1–2), Article 1–2. 10.1002/jnr.23858

Reyes-Guzman, C. M., Pfeiffer, R. M., Lubin, J., Freedman, N. D., Cleary, S. D., Levine, P. H., & Caporaso, N. E. (2017). Determinants of Light and Intermittent Smoking in the United States: Results from Three Pooled National Health Surveys. Cancer Epidemiology, Biomarkers & Prevention: A Publication of the American Association for Cancer Research, Cosponsored by the American Society of Preventive Oncology, 26(2), Article 2. 10.1158/1055-9965.EPI-16-0028

Risner, M. E., & Goldberg, S. R. (1983). A comparison of nicotine and cocaine self-administration in the dog: Fixed-ratio and progressive-ratio schedules of intravenous drug infusion. The Journal of Pharmacology and Experimental Therapeutics, 224(2), Article 2.

Samaha, A.-N., Khoo, S. Y.-S., Ferrario, C. R., & Robinson, T. E. (2021). Dopamine “ups and downs” in addiction revisited. Trends in Neurosciences, 44(7), Article 7. 10.1016/j.tins.2021.03.003

Samaha, A.-N., Yau, W.-Y. W., Yang, P., & Robinson, T. E. (2005). Rapid delivery of nicotine promotes behavioral sensitization and alters its neurobiological impact. Biological Psychiatry, 57(4), Article 4. 10.1016/j.biopsych.2004.11.040

Sanchez, V., Moore, C. F., Brunzell, D. H., & Lynch, W. J. (2014). Sex differences in the effect of wheel running on subsequent nicotine-seeking in a rat adolescent-onset self-administration model. Psychopharmacology, 231(8), Article 8. 10.1007/s00213-013-3359-3

Sannerud, C. A., Prada, J., Goldberg, D. M., & Goldberg, S. R. (1994). The effects of sertraline on nicotine self-administration and food-maintained responding in squirrel monkeys. European Journal of Pharmacology, 271(2–3), Article 2–3. 10.1016/0014-2999(94)90807-9

Schechter, M. D., & Jellinek, P. (1975). Evidence for a cortical locus for the stimulus effect of nicotine. European Journal of Pharmacology, 34(1), Article 1. 10.1016/0014-2999(75)90226-5

Shiffman, S., Dunbar, M. S., Li, X., Scholl, S. M., Tindle, H. A., Anderson, S. J., & Ferguson, S. G. (2014). Smoking Patterns and Stimulus Control in Intermittent and Daily Smokers. PLoS ONE, 9(3), Article 3. 10.1371/journal.pone.0089911

Shiffman, S., Scholl, S. M., Mao, J., Ferguson, S. G., Hedeker, D., Primack, B., & Tindle, H. A. (2020). Using Nicotine Gum to Assist Nondaily Smokers in Quitting: A Randomized Clinical Trial - PubMed. Nicotine & Tobacco Research: Official Journal of the Society for Research on Nicotine and Tobacco, 22(3), 390–397. 10.1093/ntr/ntz090

Singer, B. F., Fadanelli, M., Kawa, A. B., & Robinson, T. E. (2018). Are Cocaine-Seeking “Habits” Necessary for the Development of Addiction-Like Behavior in Rats? The Journal of Neuroscience: The Official Journal of the Society for Neuroscience, 38(1), Article 1. 10.1523/JNEUROSCI.2458-17.2017

Sorge, R. E., & Clarke, P. B. S. (2009). Rats self-administer intravenous nicotine delivered in a novel smoking-relevant procedure: Effects of dopamine antagonists. The Journal of Pharmacology and Experimental Therapeutics, 330(2), Article 2. 10.1124/jpet.109.154641

Stennett, A., Krebs, N. M., Liao, J., Richie, J. P., & Muscat, J. E. (2018). Ecological momentary assessment of smoking behaviors in native and converted intermittent smokers. The American Journal on Addictions, 27(2), Article 2. 10.1111/ajad.12690

Swalve, N., Smethells, J. R., & Carroll, M. E. (2016). Sex Differences in the Acquisition and Maintenance of Cocaine and Nicotine Self-Administration in Rats. Psychopharmacology, 233(6), Article 6. 10.1007/s00213-015-4183-8

Tapia, M. A., Jin, X.-T., Tucker, B. R., Thomas, L. N., Walker, N. B., Kim, V. J., Albertson, S. E., Damuka, N., Krizan, I., Edassery, S., Savas, J. N., Solingapuram Sai, K. K., Jones, S. R., & Drenan, R. M. (2022). Relapse-like behavior and nAChR sensitization following intermittent access nicotine self-administration. Neuropharmacology, 212, 109066. 10.1016/j.neuropharm.2022.109066

Tobacco. World Health Organisation. 2020. URL: https://www.who.int/en/news-room/fact-sheets/detail/tobacco [accessed 2021-01-05].

Watkins, S. S., Epping-Jordan, M. P., Koob, G. F., & Markou, A. (1999). Blockade of nicotine self-administration with nicotinic antagonists in rats. Pharmacology, Biochemistry, and Behavior, 62(4), Article 4. 10.1016/s0091-3057(98)00226-3

Zhai, D., van Stiphout, R., Schiavone, G., De Raedt, W., & Van Hoof, C. (2022). Characterizing and Modeling Smoking Behavior Using Automatic Smoking Event Detection and Mobile Surveys in Naturalistic Environments: Observational Study. JMIR mHealth and uHealth, 10(2), Article 2. 10.2196/28159

Zimmer, B. A., Dobrin, C. V., & Roberts, D. C. S. (2011). Brain-cocaine concentrations determine the dose self-administered by rats on a novel behaviorally dependent dosing schedule. Neuropsychopharmacology: Official Publication of the American College of Neuropsychopharmacology, 36(13), Article 13. 10.1038/npp.2011.165

Zimmer, B. A., Oleson, E. B., & Roberts, D. C. (2012). The Motivation to Self-Administer is Increased After a History of Spiking Brain Levels of Cocaine. Neuropsychopharmacology, 37(8), Article 8. 10.1038/npp.2012.37

