## Supplemental methods for "Intermittent Nicotine Access is as Effective as Continuous Access in Promoting Nicotine Seeking and Taking in Rats"

***Supplemental Information***

### Supplemental Methods

Subjects

Female (150-175 g; n = 16) and male (225-250 g; n = 16) Wistar rats (Charles River Laboratories, Saint Constant, QC) were housed individually in a temperature-controlled room (21°C ± 2°C) under a reverse 12h/12h dark/light cycle (lights off at 8:30 am). Experiments were conducted during the dark phase of the animals' circadian cycle. Water was available ad libitum and food (Charles River standard rat chow) was restricted to 20 g/day for females and 25 g/day for males, as we have used previously (Algallal et al., 2020). Mild food restriction produces healthier rats compared to ad libitum feeding, which promotes excessive fat deposition and obesity (Martin et al., 2010; Rowland, 2007).

Apparatus

Rats were trained and tested in standard operant cages (Med Associates, St Albans, VT) located in a testing room separate from the rats' housing room. Each operant cage was equipped with two retractable levers. Pressing the active lever produced reinforcement (food pellet or intravenous nicotine), pressing the inactive lever had no programmed consequences. The side of the active vs. inactive lever was kept consistent for each rat until the end of the experiment. To signal the beginning of each session, both levers were inserted into the cage and the house light was illuminated. Upon reinforcer delivery and for the ensuing 20-s timeout period (25 s total), both levers were retracted, the light above the active lever was illuminated, and a 2900-Hz, 75-dB tone was presented for the first 5 s. After the 25 s, the light above the active lever was extinguished, and the levers were inserted again to indicate reinforcer availability. This light-tone stimulus was used as a nicotine-associated conditioned stimulus in later cue-induced reinstatement tests, as detailed below. After the timeout period, the levers were inserted back into the cage to indicate reward availability. A computer running Med Associates Med-PC (v. 4) software controlled testing parameters in the cages and collected data.

Food self-administration training and catheter implantation

To hasten the acquisition of lever pressing for nicotine, rats were first trained to lever-press for 45-mg, grain-based pellets (product # F0165, Bioserv; VWR), under a fixed ratio 1 schedule of reinforcement (FR1), with a 25 s timeout period following each reinforcer delivery. Sessions lasted 1 h or until 100 pellets were self-administered. Following two training sessions, one female and one male rat did not meet the acquisition criterion of earning at least ~25 pellets/session and were given an overnight session. Once rats met the criterion, the schedule of reinforcement was changed to FR3 for at least 2 sessions.

On the day following acquisition of reliable operant responding for food (as indicated by the self-administration of ~25 pellets/session, on two consecutive sessions), the rats were anesthetized with isoflurane (5% for induction, 2-3% for maintenance) and catheters were implanted into the right jugular vein, as in previous work (Samaha et al., 2011; Weeks, 1962). Two female rats died during surgery. On the day of surgery, rats received an intramuscular injection of a penicillin antibiotic (Procillin; CDMV, St Hyacinthe, Qc) and a subcutaneous injection of the anti-inflammatory agent Carprofen (Rimadyl; CDMV, St Hyacinthe, Qc). Thereafter, catheters were flushed on alternate days with saline and saline containing 0.2 mg/ml of heparin (Sigma-Aldrich, Oakville, ON), and 2 mg/ml of Baytril (CDMV, St Hyacinthe, QC).

Drugs

During nicotine self-administration sessions, nicotine tartrate (MP Biomedicals; 0.015 mg/kg/injection, calculated as the weight of the base) was dissolved in 0.9% saline with pH adjusted to 7.2-7.4 and injected over 5 seconds (31 µl/s) using syringe-pumps with 3.33-RPM motors (Med Associates; model no. PHM-100).

Nicotine self-administration training

Following recovery from surgery (5-7 days), rats learned to self-administer nicotine (0.015 mg/kg/injection, delivered over 5 s, followed by a 20-s timeout period) under FR3, during 1-h sessions, 1 session/day. Immediately before each session, to enable rats to move freely, their catheters were connected to a nicotine-containing syringe via tubing protected by a stainless-steel spring connected to a liquid swivel set in a counterbalanced arm. Rats were given daily sessions until the following acquisition criteria were met on two consecutive sessions; taking ≥ 6 injections/session, pressing at least twice more on the active versus inactive lever, and taking nicotine in a regular pattern throughout the session, as indicated by visual inspection of cumulative response records. One female rat was excluded for not meeting these criteria, and 4 females and 3 males were excluded because of loss of catheter patency. In addition, one female rat was excluded because of aggressive behaviour and another female rat was excluded because problems with her teeth hindered normal feeding.

Assessing catheter patency

Catheter patency was verified after the last LgA/IntA nicotine self-administration session and at the end of progressive ratio testing. At each time point, rats were given an i.v. infusion of propofol (10 mg/mL; 0.1 mL/rat; CDMV, St-Hyacinthe, QC), a short-acting anaesthetic [$T_{1/2}$ ~27 minutes in Wistar rats; (Dutta et al., 1997)]. Rats that did not become ataxic within 10 s of the infusion were considered to have non-patent catheters and were excluded from data analysis.

Modeling brain nicotine concentrations

Central nicotine concentrations were estimated using self-administration time series data from the 12th LgA and IntA sessions. For pharmacokinetic modeling, we used Tapia et al's (2022) approach, using the same parameters and the MATLAB/SimBiology code downloaded from their github repository (version uploaded on Dec 2nd, 2021) and provided with the original paper. Briefly, the pharmacokinetic model included two compartments. The central compartment corresponded to blood, plasma, and cerebrospinal fluid. The peripheral compartment included fat, muscle, and other poorly perfused tissues. The time evolution of simulated nicotine concentration in the central compartment (hereafter referred to as plasma nicotine concentration) was used in subsequent analyses. The model was validated in male rats (Tapia et al., 2022) by comparing central nicotine kinetics predicted by the model to rat plasma nicotine kinetics after arterial bolus injection [(Kyerematen et al., 1988) see also (Craig et al., 2014)]. Because parameter fitting was based on male rat data, pharmacokinetic modeling was applied only to male rats in each group. From the time course of the (simulated) plasma nicotine concentrations, we identified Cmax (in ng/mL) and Tmax (in minutes) values. Moreover, we computed area under the curve (AUC; in ng/mL * h) from the beginning of the session to different time points thereafter (2-24 h).

Data analysis

Estimated nicotine Tmax and Cmax values were compared between groups (LgA or IntA) using unpaired *t* tests. We used two-way repeated-measures ANOVA to analyze area under the curve (Access x Time; Time as a within-subjects variable). Lever pressing behavior during the 6-h sessions was analysed with three-way repeated-measures ANOVA (Access (LgA or IntA) x Lever type x Session; the latter two as within-subjects variables). We used two-way repeated-measures ANOVA to analyse the number of self-administered infusions during the 6-h sessions (Access x Session; Session as a within-subjects variable) and during PR tests (Access x Dose; Dose as a within-subjects variable). Group differences in active lever pressing rates during

LgA/IntA sessions and cumulative nicotine intake after these sessions were analyzed using unpaired *t* tests. We also used three-way repeated-measures ANOVA to analyze lever-pressing behaviour during extinction sessions (Access x Time x Lever type; the latter two as within-subjects variables) and CS- and nicotine-induced reinstatement tests (Access x Test type (Extinction vs. Reinstatement) x Lever type; the latter two as within-subjects variables). When data violated sphericity, a Greenhouse-Geisser correction was applied or Welch's t-test was conducted, as appropriate. To assess the relationship between measures of responding during LgA/IntA sessions on the one hand (i.e., cumulative intake, inter-infusion interval and lever pressing rate) and progressive ratio or reinstatement responding on the other, we analyzed goodness-of-fit ($r^2$) of the linear regression between these variables. Alpha level was set at $p \leq 0.05$, except for analysis of the significance of correlations, where we applied Bonferroni's correction to adjust for multiple comparisons. Data were analyzed with GraphPad Prism (v. 7.0d) and SPSS (v. 20). Data in figures are mean ± SEM.

**References**

Algallal, H., Allain, F., Ndiaye, N. A., & Samaha, A.-N. (2020). Sex differences in cocaine self-administration behaviour under long access versus intermittent access conditions. *Addiction Biology*, *25*(5), Article 5. https://doi.org/10.1111/adb.12809

Craig, E. L., Zhao, B., Cui, J. Z., Novalen, M., Miksys, S., & Tyndale, R. F. (2014). Nicotine pharmacokinetics in rats is altered as a function of age, impacting the interpretation of animal model data. *Drug Metabolism and Disposition: The Biological Fate of Chemicals*, *42*(9), Article 9. https://doi.org/10.1124/dmd.114.058719

Dutta, S., Matsumoto, Y., & Ebling, W. F. (1997). Propofol pharmacokinetics and pharmacodynamics assessed from a cremophor EL formulation. *Journal of Pharmaceutical Sciences*, *86*(8), Article 8. https://doi.org/10.1021/js970118m

Kyerematen, G. A., Taylor, L. H., deBethizy, J. D., & Vesell, E. S. (1988). Pharmacokinetics of nicotine and 12 metabolites in the rat. Application of a new radiometric high performance liquid chromatography assay. *Drug Metabolism and Disposition: The Biological Fate of Chemicals*, *16*(1), Article 1.

Martin, B., Ji, S., Maudsley, S., & Mattson, M. P. (2010). “Control” laboratory rodents are metabolically morbid: Why it matters. *Proceedings of the National Academy of Sciences of the United States of America*, *107*(14), Article 14. https://doi.org/10.1073/pnas.0912955107

Rowland, N. E. (2007). Food or fluid restriction in common laboratory animals: Balancing welfare considerations with scientific inquiry. *Comparative Medicine*, *57*(2), Article 2.

Samaha, A.-N., Minogianis, E.-A., & Nachar, W. (2011). Cues paired with either rapid or slower self-administered cocaine injections acquire similar conditioned rewarding properties. *PloS One*, *6*(10), Article 10. https://doi.org/10.1371/journal.pone.0026481

Tapia, M. A., Jin, X.-T., Tucker, B. R., Thomas, L. N., Walker, N. B., Kim, V. J., Albertson, S. E., Damuka, N., Krizan, I., Edassery, S., Savas, J. N., Solingapuram Sai, K. K., Jones, S. R., & Drenan, R. M. (2022). Relapse-like behavior and nAChR sensitization following intermittent access nicotine self-administration. *Neuropharmacology*, *212*, 109066. https://doi.org/10.1016/j.neuropharm.2022.109066

Weeks, J. R. (1962). Experimental morphine addiction: Method for automatic intravenous injections in unrestrained rats. *Science (New York, N.Y.)*, *138*(3537), Article 3537. https://doi.org/10.1126/science.138.3537.143
