## Supplemental figure 1 for "Intermittent Nicotine Access is as Effective as Continuous Access in Promoting Nicotine Seeking and Taking in Rats"

**Supplementary Figure 1**

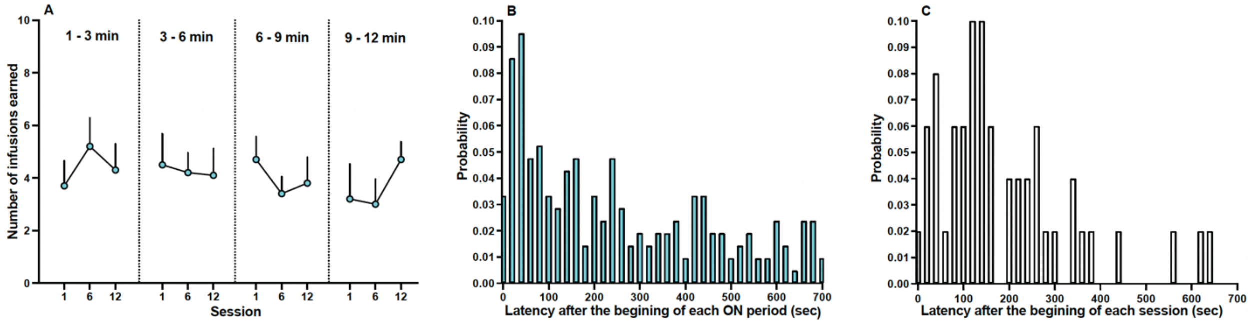


Analysis of the temporal pattern of nicotine intake in IntA rats. (**A**) During the 12-min nicotine ON periods of sessions 1, 6 and 12, IntA rats earned comparable numbers of nicotine infusions in minutes 1-3, minutes 3-6, minutes 6-9, and minutes 9-12. Data are mean ± SEM. n = 10. Probabilistic analysis of the latency to self-administer a nicotine infusion after (**B**) the start of a nicotine ON period in IntA rats (**C**) the start of a LgA session, over the last five sessions (8 to 12). n = 7 females, 13 males
