## Supplemental tables for "Intermittent Nicotine Access is as Effective as Continuous Access in Promoting Nicotine Seeking and Taking in Rats"

**Supplementary table 1:**
**Correlations between each of cumulative nicotine intake, inter-infusion interval and lever pressing rate and later responding for nicotine (0.0075, 0.015 & 0.03 mg/kg/infusion) under a progressive ratio schedule of reinforcement.**

| | | Nicotine infusions earned under progressive ratio | | | | | |
|---|---|---|---|---|---|---|---|
| | | 0.0075 | | 0.015 | | 0.03 | |
| | | $r^2$ | $p$ | $r^2$ | $p$ | $r^2$ | $p$ |
| Cumulative intake | IntA | 0.264 | 0.129 | 0.246 | 0.145 | 0.320 | 0.088 |
| | LgA | 0.342 | 0.076 | 0.755 | 0.001* | 0.337 | 0.078 |
| Inter-infusion Interval | IntA | 0.196 | 0.200 | 0.255 | 0.137 | 0.004 | 0.867 |
| | LgA | 0.155 | 0.260 | 0.037 | 0.592 | 0.266 | 0.127 |
| Lever pressing/min | IntA | 0.322 | 0.870 | 0.142 | 0.283 | 0.117 | 0.334 |
| | LgA | 0.214 | 0.178 | 0.672 | 0.004 | 0.182 | 0.219 |

**Supplementary table 2:**
**Correlations between each of cumulative nicotine intake, inter-infusion interval and lever pressing rate and active lever pressing during later cue- and nicotine-induced reinstatement tests.**

| | | Cue-induced Reinstatement | | Nicotine-induced Reinstatement | |
|---|---|---|---|---|---|
| | | $r^2$ | $p$ | $r^2$ | $p$ |
| Cumulative intake | IntA | 0.003 | 0.870 | 0.185 | 0.214 |
| | LgA | 0.094 | 0.389 | 0.048 | 0.542 |
| Inter-infusion Interval | IntA | 0.275 | 0.118 | 0.018 | 0.709 |
| | LgA | 0.639 | 0.005 | 0.092 | 0.395 |
| Lever pressing/min | IntA | 0.150 | 0.268 | 0.102 | 0.369 |
| | LgA | 0.022 | 0.688 | 0.07 | 0.455 |
